# The Gehan test identifies life-extending compounds overlooked by the log-rank test in the NIA Interventions Testing Program: Metformin, Enalapril, caffeic acid phenethyl ester, green tea extract, and 17-dimethylaminoethylamino-17-demethoxygeldanamycin hydrochloride

**DOI:** 10.1101/2024.02.17.579363

**Authors:** Nisi Jiang, Jonathan Gelfond, Qianqian Liu, Randy Strong, James F. Nelson

## Abstract

The National Institute on Aging Interventions Testing Program (ITP) has so far identified 12 compounds that extend the lifespan of genetically heterogeneous mice using the log-rank test. However, the log-rank test is relatively insensitive to any compound that does not uniformly reduce mortality across the lifespan. This test may thus miss compounds that only reduce mortality before midlife, for example, a plausible outcome if a compound only mitigates risk factors before midlife or if its efficacy is reduced at later ages. We therefore reanalyzed all data collected by the ITP from 2004-2022 using the Gehan test, which is more sensitive to mortality differences earlier in the life course and does not assume a uniformly reduced mortality hazard across the lifespan. The Gehan test identified 5 additional compounds, metformin, enalapril, 17-dimethylaminoethylamino-17-demethoxygeldanamycin hydrochloride (17-DMAG), caffeic acid phenethyl ester (CAPE), and green tea extract (GTE), which significantly increased survival but were previously missed by the log-rank test. Three (metformin, enalapril, and 17-DMAG) were only effective in males and two (CAPE and GTE) were only effective in females. In addition, 1,3-butanediol, which by log-rank analysis increased survival in females but not males, increased survival in males by the Gehan test. These results suggest that statistical tests sensitive to non-uniformity of drug efficacy across the lifespan should be included in the standard statistical testing protocol to minimize overlooking geroprotective interventions.

## Introduction

The National Institute on Aging Interventions Testing Program (ITP) aims to identify drugs and other compounds that, when provided in the food of genetically heterogeneous mice, are broadly geroprotective [1]. Extension of population survival is the initial criterion that must be met for further testing on other measures of aging, such as frailty, sarcopenia, and cognitive decline. To determine the life-extending effect of a compound, the ITP uses the log-rank test, which assumes an effect on mortality hazard independent of age [2], combined with the Allison-Wang test to assess effects on maximum lifespan [3]. Using these tests, twelve of the forty-eight drugs tested, including notable examples like rapamycin and acarbose, have had positive impacts on lifespan [4-16].

Aging research typically focuses on the later ages of life. However, a well-accepted measure of aging, the exponential increase of mortality with advancing chronological age, begins in early adulthood [17]. Since administration of most interventions tested in ITP begins early in adult life, it is possible that some might only influence mortality in early adulthood or middle age but have minimal effects at later ages. For example, prepubertal castration, a non-pharmacological intervention, reduces male mortality up to the median age of survival but has no effect thereafter [18]. Although interventions that only attenuate mortality during the first half of life are not geroprotective during the senescent stage of life, they are therapeutically consequential for a substantial period of life. In addition, these treatments occur during a period when the exponentially increasing mortality hazard has already begun. Their efficacy, limited to the first half of life, could reflect action on pathways that mainly cause aging in early and mid-adulthood. Indeed, human epidemiological data shows that causes of death vary with age, suggesting that the underlying biology responsible for mortality may vary across the life course [19]. Alternatively, age-related alterations in pharmacokinetics and pharmacodynamics might underlie their reduced efficacy at later ages, a result that might require modifications to extend their efficacy to later ages [20].

The log-rank test lacks sensitivity for identifying potential effects on lifespan limited to early stages of life because it weights the differences in mortality hazard across all ages equally and assumes the hazard functions of the groups being compared are proportional (proportional hazard (PH) assumption) [21]. These assumptions make the log-rank test most sensitive to interventions with consistent effects on mortality through the lifespan. However, the test is less suitable for identifying interventions with effects during a limited age period, since the relative mortality hazard risk between groups will change over time and thus will no longer fit the PH assumption.

To increase the likelihood of identifying interventions that influence lifespan mainly during the first half of the life course, we reevaluated the ITP survival data using an alternative statistical analysis, the Gehan-Breslow-Wilcoxon (Gehan) test [22, 23]. The Gehan test is calculated in a similar way as the log-rank test, but unlike the log-rank test, it does not assume proportional hazards. The Gehan test is thus preferable to the log-rank test for interventions that have effects only during certain ages and do not fit the PH assumption. In addition, the Gehan test weights different ages proportionate to the number of individuals still alive, which makes the Gehan test more sensitive to the interventions that have age-specific effects, especially during the early ages [24]. Of note, this analysis has been used in the ITP several times to determine the potential drug effects [6, 7], but never systematically. Using this approach, we have found 6 additional interventions that have significant effects on lifespan, particularly in the early portion of life.

## Results

We first re-evaluated all the available published survival data of compounds tested by the ITP to determine if they met the assumption of the log-rank test that the proportional hazard was equal across the entire age range of treatment. Of 132 cases (drugs/conditions/sexes/cohorts), 16 did not meet this assumption, making the log-rank test less sensitive for these cases (Table S1 and S2). Of these 16 compounds, the results using the log-rank and Gehan tests were consistent for 13 cases (Table S1). However, for the other 3 compounds, metformin and enalapril in males, and Green Tea Extract (GTE) in females, which were not significant by log-rank analysis, their life-extending effect was significant using the Gehan test (Table 1). The remaining 116 cases met the PH assumption and thus both the log-rank and Gehan tests are appropriate statistically. Within this group, the Gehan test identified 3 additional compounds that increased survival but had not been identified previously by the log-rank Test: caffeic acid phenethyl ester (CAPE) in females, 17-dimethylaminoethylamino-17-demethoxygeldanamycin hydrochloride (17-DMAG) in males, and 1,3-butanediol (BD) in males (Table 1). It should be noted that BD was originally found by the log-rank test to be significant in females but not in males. Figure 1 shows the Kaplan-Meier survival plots for these newly identified life-extending compounds.

**Table 1.** Statistical results for log-rank, Gehan, and Cox ZPH tests in drugs that were newly identified as lifespan-extending. The log-rank and Gehan tests are described in the Methods. The Z-test evaluates if the hazard ratio between drug-treated, and control mice is constant throughout the treatment period and adheres to the proportional hazard assumption. If violated, the Gehan test could be used.

| Treatment | Sex | Dosage (ppm) | Age of onset (month) | Median lifespan changes | Log_rank p_value | Gehan p_value | Ztest p_value | PH<sup>1</sup> Assumption |
| --- | --- | --- | --- | --- | --- | --- | --- | --- |
| GTE | f | 2000 | 4 | +7% | 0.3686 | 0.0344 | 0.0023 | Violated |
| Met | m | 1000 | 9 | +8% | 0.3477 | 0.0495 | 0.0118 | Violated |
| Enal | m | 120 | 4 | +7% | 0.2200 | 0.0460 | 0.0343 | Violated |
| CAPE | f | 300 | 4 | +5% | 0.0665 | 0.0481 | 0.4660 | Not Violated |
| DMAG | m | 30 | 6 | +7% | 0.1173 | 0.0321 | 0.1486 | Not Violated |
| BD | m | 100000 | 6 | +9% | 0.1097 | 0.0248 | 0.0533 | Not Violated |

<sup>1</sup> Proportional Hazard

**Fig. 1.**
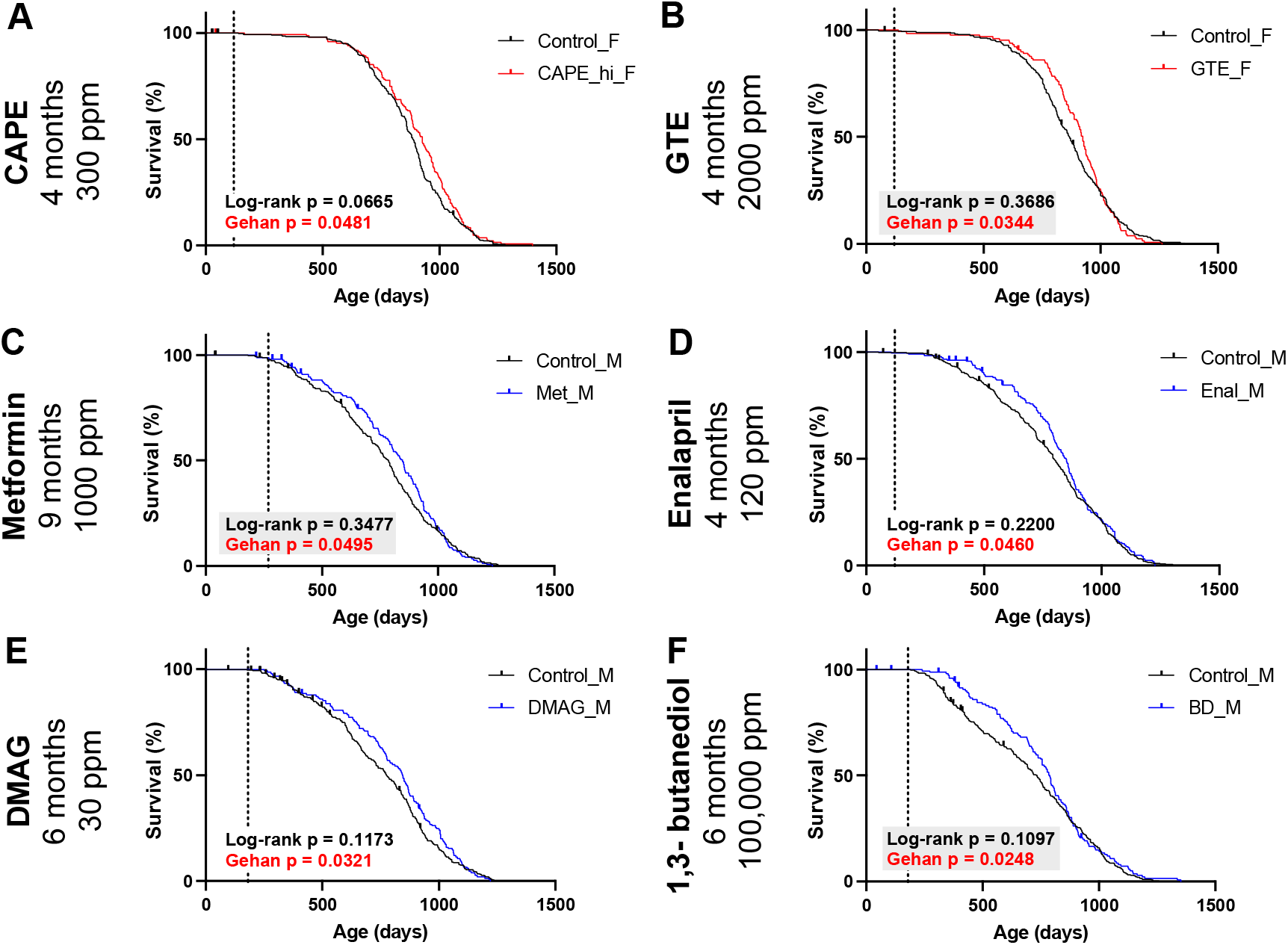
Survival curves and statistics for drugs that significantly increase survival according to the Gehan test, but do not do so according to the log-rank test. (The vertical dotted line is the age of treatment onset.) **(A)** Caffeic Acid Phenethyl Ester treated (CAPE, 300 ppm, from 4-month-old, n = 147) and control (n = 289) female mice. **(B)** Green Tea Extract treated (GTE, 2000 ppm, from 4-month-old, n = 129) and control (n = 275) female mice. **(C)** Metformin treated (1000 ppm, from 9-month-old, n = 148) and control (n = 294) male mice. **(D)** Enalapril treated (120 ppm, from 4-month-old, n = 170) and control (n = 357) male mice. **(E)** 17-dimethylaminoethylamino-17-demethoxygeldanamycin hydrochloride treated (17-DMAG, 30 ppm, from 6-month-old, n = 152) and control (n = 264) male mice. **(F)** 1,3-butanediol treated (BD, 100,000 ppm, from 6-month-old, n = 145) and control (n = 285) male mice.

### Caffeic Acid Phenethyl Ester prolonged lifespan in females

CAPE was selected for ITP because of its potent anti-inflammatory, antioxidant, and anticancer properties, and was initially tested by the ITP at 2 doses, 30 ppm, and 300 ppm [5, 25]. However, neither dose of CAPE increased lifespan significantly using the log-rank test. In our analysis, 300 ppm CAPE-treated female mice lived significantly (p=0.0481) longer than the control (Figure 1A). The median lifespan of CAPE-treated females (926 days) is 5% longer than that of controls (886 days). However, the 300-ppm treated male group remained non-significant (Figure S1A).

### Green Tea Extract prolonged lifespan in females

GTE was proposed to ITP due to its strong antioxidant properties and tested at 2000 ppm from 4 months of age onward [7]. Although it prolonged the female median lifespan by 7%, this was not significant by log-rank testing [7]. In our analysis, GTE-treated females violated the PH assumption, possibly because the difference in survival between the GTE-treated group and the control group greatly diminished in advanced age (Figure 1B). Using the Gehan test, GTE increased survival significantly in females (p=0.0344), but not in males (Figure S1B). Of note, the Gehan test was used as a secondary analysis in the original report and showed a similar outcome [7]. It is worth noting that the progressively diminishing effect on survival was also observed in an earlier study using B6 mice, where the average lifespan was significantly longer in GTE-treated mice, but there was no effect on maximum lifespan [26]. The diminished effect of GTE at later ages suggests that GTE may have ameliorated specific risk factors that are non-uniform across the lifespan.

### Metformin prolonged lifespan in males

Metformin at 1000 ppm starting at 9 months increased the median lifespan of males by 8% (Figure 1C), but this effect did not reach statistical significance using the log-rank test [9]. Here we found that metformin treatment of males violated the PH assumption (Table 1), indicating the effect of metformin on mortality changed during age. Analysis using the Gehan test showed a significant increase in survival of metformin-treated males (p=0.0495), but not in females (Figure 1C and S1C).

### Enalapril prolonged lifespan in males

Enalapril is an angiotensin-converting enzyme (ACE) inhibitor. Treatment at 120 ppm beginning at 4 months prolonged median lifespan by 7% in males (Figure 1D), but not significantly using the log-rank test [5]. Here we found that enalapril treatment of males violated the PH assumption (Table 1), suggesting a significant decline in the effect of enalapril on mortality changes across the lifespan. Analysis using the Gehan test showed a significant increase in survival of enalapril-treated males (p=0.0343), but not in females (Figure 1D and S1D).

### 17-dimethylaminoethylamino-17-demethoxygeldanamycin hydrochloride prolonged lifespan in males

Treatment with 30 ppm of 17-DMAG, an HSP90 inhibitor [27], beginning at 6 months prolonged median lifespan by 7% in males (Figure 1E), but not significantly using the log-rank test [12]. Although its effect on mortality appears diminished in the advanced age (Figure 1E), it did not significantly violate the PH assumption (Table 1). In this case, both Gehan and log-rank tests are applicable. Analysis using the Gehan test showed that 17-DMAG significantly (p=0.0321) increased survival of males, but not in females (Figure S1E). Since this effect is only significant in the Gehan test and not the log-rank test, it suggests the efficacy of the drug may be mainly limited to earlier ages.

### 1,3-butanediol (BD) prolonged lifespan in males

BD was tested because of its ketogenic effect, and treatment at 100,000 ppm, beginning at 6 months, increased the median lifespan of males by 9% (Figure 1F), but not significantly using the log-rank test. By contrast, in females, the median lifespan increased by only 2% (Figure S1F) but reached statistical significance by log-rank testing [15]. Although the effect on survival in both sexes appeared to diminish with advancing age, BD treatment in both sexes did not violate the PH assumption (Table 1), making both the log-rank and Gehan tests applicable. Using the Gehan test, the effect of BD on males was significant (p=0.0248) while the effect on females became nonsignificant (p=0.1632).

### Three log-rank-identified drugs lost their significance in the Gehan test

Three drugs that had been identified to extend lifespan using the log-rank test no longer had statistically significant effects according to the Gehan test: acarbose in females at 400 ppm and 1000 ppm (Figure S2A and S2B), butanediol in females (Figure S2C), and glycine in males (Figure S2D). Acarbose has been tested in 10 trials with varying doses and durations. In all other trials except those in females at 400 and 1000 ppm, the Gehan and log-rank tests are consistent in terms of assigning significance to the effect (Table S2). None of these drugs at the doses noted violated the PH assumption. Thus, both Gehan and log-rank tests can be used. The loss of significance using the Gehan test may indicate the drug effects are stronger at later ages when the Gehan test loses sensitivity.

## Discussion

A major goal of the ITP is to identify geroprotective candidates deserving of follow-up and confirmation, and increasingly the ITP is looked to as the most robust and reliable indicator of such candidates. Thus, it is especially important that the experimental design of the ITP minimizes false negatives, which are unlikely to be pursued further. We undertook this re-evaluation because visual examinations of the Kaplan-Meier survival plots of the compounds tested by the ITP suggested that the efficacy of many of the compounds appeared to vary with age, often appearing to be less effective as the maximum lifespan was approached (Figure 1). The log-rank test is relatively insensitive to interventions whose efficacy is not uniform across the lifespan and thus may not identify such agents [21]. We chose the Gehan test for this reevaluation as a complement to the log-rank test because the Gehan test does not require the proportional hazard to be invariant across the testing age range. Moreover, it has a gradient of sensitivity that is greatest at the outset of the testing interval, declining thereafter as the size of the surviving population progressively declines, making it especially suitable for those interventions that appeared to lose efficacy at older ages.

Reevaluation of the ITP dataset using the Gehan test identified five compounds that had not been reported as geroprotective based on the standard statistical testing protocol of the Interventions Testing Program. Also, the Gehan test identified one compound to be effective in males that had only been identified by log-rank testing as effective in females. This represents a nearly 50% increase in the number of compounds identified by the ITP that may enhance survival at least during part of the adult lifespan.

In our reanalysis, metformin, and enalapril have emerged as candidates warranting further exploration, especially given their wide use clinically. Metformin is the first-line FDA-approved antidiabetic drug and has been widely used in humans for the treatment of type 2 diabetes [28]. It has been shown to prolong lifespan in C. elegans and inbred male mice [29, 30], and reduce all-cause mortality in humans [31]. The initial report of the ITP [9], in which metformin increased the median lifespan of males by 8% albeit not significantly by the log-rank test, was a marked contrast to these observations. Our finding of a significant effect of metformin on lifespan by the Gehan test suggests that metformin may be more impactful on survival at earlier ages with little effect in the late stage of life. It is noteworthy that the earlier study of metformin in C57BL/6 and B6C3F1 mice also used the Gehan test and showed a similar Kaplan-Meier survival plot with diminished efficacy at later ages [30].

Enalapril, an ACE inhibitor widely used in humans for blood pressure management, was selected for testing because of its beneficial effects on hypertension, obesity, diabetes, and congestive heart failure in aged humans [32-34]. It has been reported to prolong lifespan in rats [35]. The observation that the beneficial effect of enalapril on lifespan can be only detected in the Gehan test suggests that, as with metformin, its beneficial effects on mortality are greatest at midlife. Of note, captopril, another ACE inhibitor tested by the ITP in a later trial, prolonged life in males according to log-rank analysis [15]. Similar to Enalapril, the mortality-reducing effects of captopril are markedly diminished with advancing age (Figure S3). These results suggest that the beneficial effects of captopril and enalapril, as well as GTE and butanediol, are limited to midlife and may not be effective at older ages.

An important outcome of this re-evaluation is the evidence it provides that a number of the compounds tested by the ITP do not appear to have a uniformly salutary effect on mortality reduction across the lifespan. The compounds newly detected by the Gehan test have relatively small effect sizes, and their effect sizes diminished in advanced ages (Figure 1). This decline could indicate a loss of drug efficacy or increase in toxicity, due to age-related alterations in drug metabolism. Drug pharmacokinetics can vary markedly with age [20], as exemplified by the age-related change in plasma levels of canagliflozin, a drug found by the ITP to reduce mortality markedly [36]. Canagliflozin levels increased nearly 4-fold in older mice given the same dose as young mice [13].

Alternatively, some interventions may only mitigate risk factors that are limited to specific ages. An example is castration, a non-pharmacological intervention that markedly reduces the mortality hazard of males, but only from puberty to the median age of survival, with no effect thereafter [18]. In this example the risk factors that castration mitigates either diminish or become insignificant in the context of greater risk factors that emerge at later ages. Another implication of this reanalysis is that more attention should be given to determining how aging affects the pharmacokinetics and pharmacodynamics of the compounds being tested, both to maximize their efficacy and to understand the aging-promoting processes they are targeting. By methodically analyzing the pharmacokinetic changes of these drugs with aging, and by designing experiments to determine whether a drug loses efficacy due to aging or merely has a time-limited impact, we can significantly advance our understanding of the mechanisms of action of these drugs as well as enhance our knowledge of the aging processes they impact.

It is noteworthy that the sex differences in drug efficacy that have characterized the ITP continue with the compounds identified here [37]. CAPE and GTE were effective only in females, notably the first drugs to be female-specific. The only other drug that has been more effective in females is rapamycin. All the other drugs have either only worked in males or been more effective in males [37]. The identification of two drugs that are female-specific may help discover the basis for the marked sex differences uncovered by the ITP. Conversely, understanding why 17-DMAG, metformin, and enalapril are male-specific can, combined with the many others previously identified, help determine if there are common targets that only affect the aging of males.

## Conclusion

Re-evaluating the ITP dataset with a statistical test that does not require invariant reduction of mortality hazard across the lifespan identified 5 new potentially geroprotective drugs and 1 drug with an effect on a sex not observed with the log-rank test. These results and the finding of interventions whose effects on the hazard ratio changed significantly across the life course suggest that the efficacy of geroprotective drugs is not always uniform across the lifespan. This finding underscores the need for intervention research to give more attention to pharmacokinetic and pharmacodynamic changes during aging and to consider life-stage-specific therapeutic interventions.

## Methods

### UM-HET3 mice and husbandry

This paper reexamines data from original reports, which detail the mouse model, breeding, drug preparation and administration, mouse observation, and data collection. UM-HET3 mice derived from BALB/cByJ x C57BL/6J F1 mothers and C3H/HeJ x DBA/2J F1 fathers. The mice were kept in conditions of 25 °C and a 12/12-hour light/dark cycle, with unrestricted access to Purina 5LG6 diet. Housing was organized with three males or five females per cage. Daily health checks were conducted to monitor for signs of illness. Mice deemed by a qualified technician to have less than 48 hours of survival, especially due to critical conditions such as severe tumors or incapacitation, were humanely euthanized. The age at euthanasia was noted as the closest approximation of the natural lifespan for these cases. For mice found deceased during routine inspections, the age at discovery was recorded as their lifespan. Instances of aggressive behavior resulting in significant injuries (bleeding, infected, or wounds covering over 20% of the body surface) led to the euthanization of all mice in the affected cage.

## Data Availability and Statistical Analysis

All survival data are acquired from the mouse phenome database (phenome.jax.org). The differences in right censored survival were tested using the log-rank and Gehan-Breslow-Wilcoxon test with the Peto modification [23]. For each gender, the ITP data was pooled from three sites with stratification by site. The log-rank test was performed in the same way as in the original publications by the Interventions Testing Program. Pooled data across the three test sites were compared with stratified by the site. The Z-test of the proportional hazard association was performed to assess the appropriate-ness of the log-rank test. To account for the multiplicity of testing (132 trials), the p-values were adjusted using a 5% FDR threshold [38]. The analysis was conducted in the R computing environment (V4.3.1, Vienna, Austria) using the *survival* R package.

## Statements and Declarations

## Acknowledgment

We thank Dr. Adam Salmon (UT Health San Antonio) for the insightful suggestions. Dr. Strong has been honored with a Senior Research Career Scientist award (# IK6 BX006289) by the Department of Veterans Affairs. Additionally, this research received support from the Center for Evaluating Potential Anti-Aging Interventions (5U01AG022307) and the Nathan Shock Center of Excellence in Basic Biology of Aging (5P30AG013319).

## Data Availability

The data and the codes that support the findings of this study are available from the corresponding author upon reasonable request.

---

**Table S1.** Comparison of log-rank, Gehan tests stratified by Cox ZPH test. The log-rank and Gehan tests are described in the Methods. The Cox ZPH test evaluates if the hazard ratio between drug treated and control mice is constant throughout the treatment period, and thus adheres to the proportional hazard assumption.

| Conditions | Total | PH Not Violated | PH Violated |
| --- | --- | --- | --- |
| Log-rank, Sig; Gehan, Sig. | 41 | 33 | 8 |
| Log-rank, Sig; Gehan, Non-Sig. | 4 | 4 | 0 |
| Log-rank, Non-Sig; Gehan, Sig. | 6 | 3 | 3 |
| Log-rank, Non-Sig; Gehan, Non-Sig. | 81 | 76 | 5 |
| Total | 132 | 116 | 16 |

**Fig. S1.**
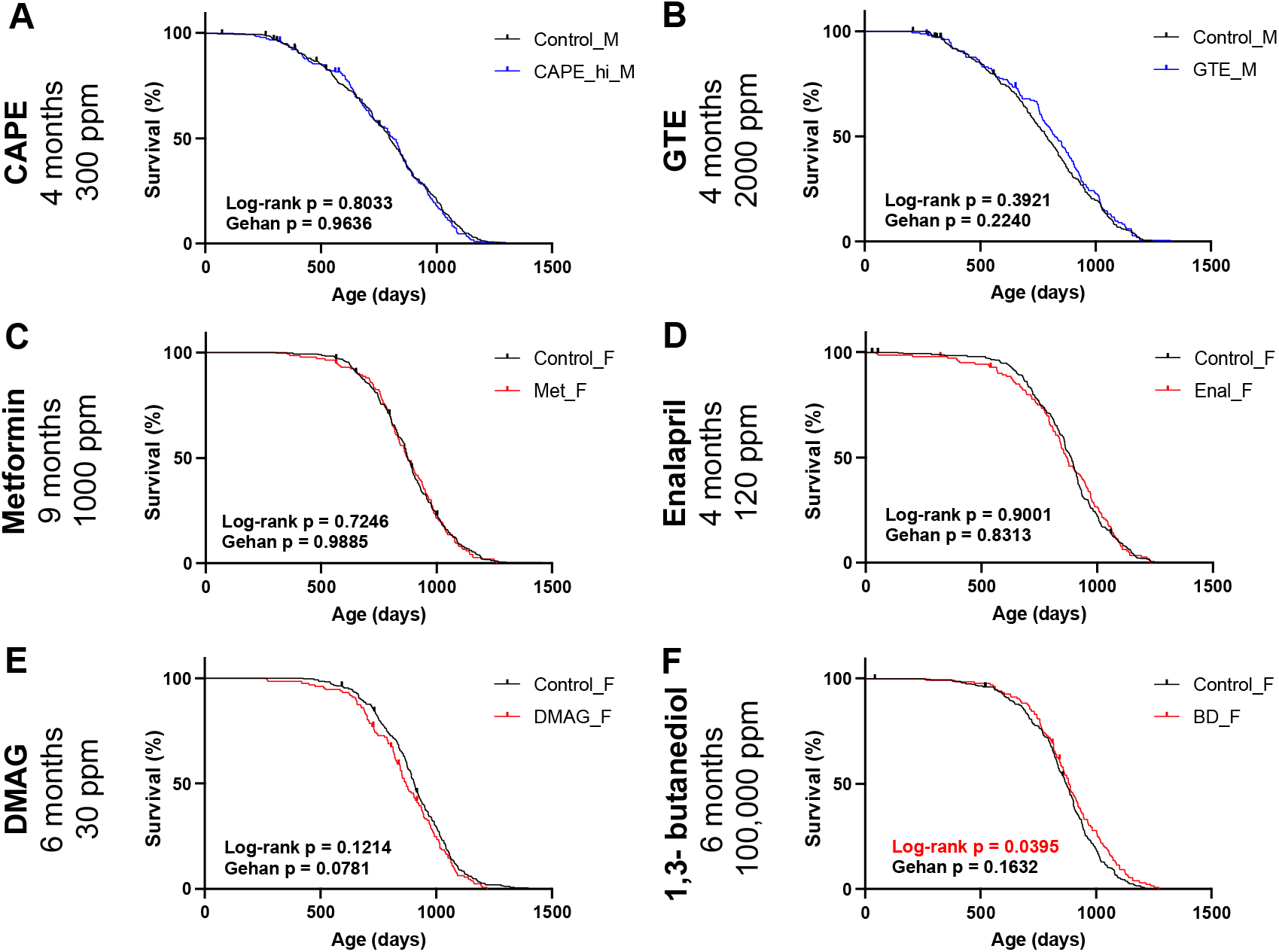
Survival curves and statistics for the other sex of the interventions newly identified by the Gehan test. **(A)** Caffeic Acid Phenethyl Ester treated (CAPE, 300 ppm, from 4-month-old, n = 171) and control (n = 375) male mice. **(B)** Green Tea Extract treated (GTE, 2000 ppm, from 4-month-old, n = 145) and control (n = 276) male mice. **(C)** Metformin treated (1000 ppm, from 9-month-old, n = 140) and control (n = 281) female mice. **(D)** Enalapril treated (120 ppm, from 4-month-old, n = 139) and control (n = 289) female mice. **(E)** 17-dimethylaminoethylamino-17-demethoxygeldanamycin hydrochloride treated (DMAG, 30 ppm, from 6-month-old, n = 132) and control (n = 274) female mice. **(F)** 1,3-butanediol treated (BD, 100,000 ppm, from 6-month-old, n = 134) and control (n = 276) female mice.

**Fig. S2.**
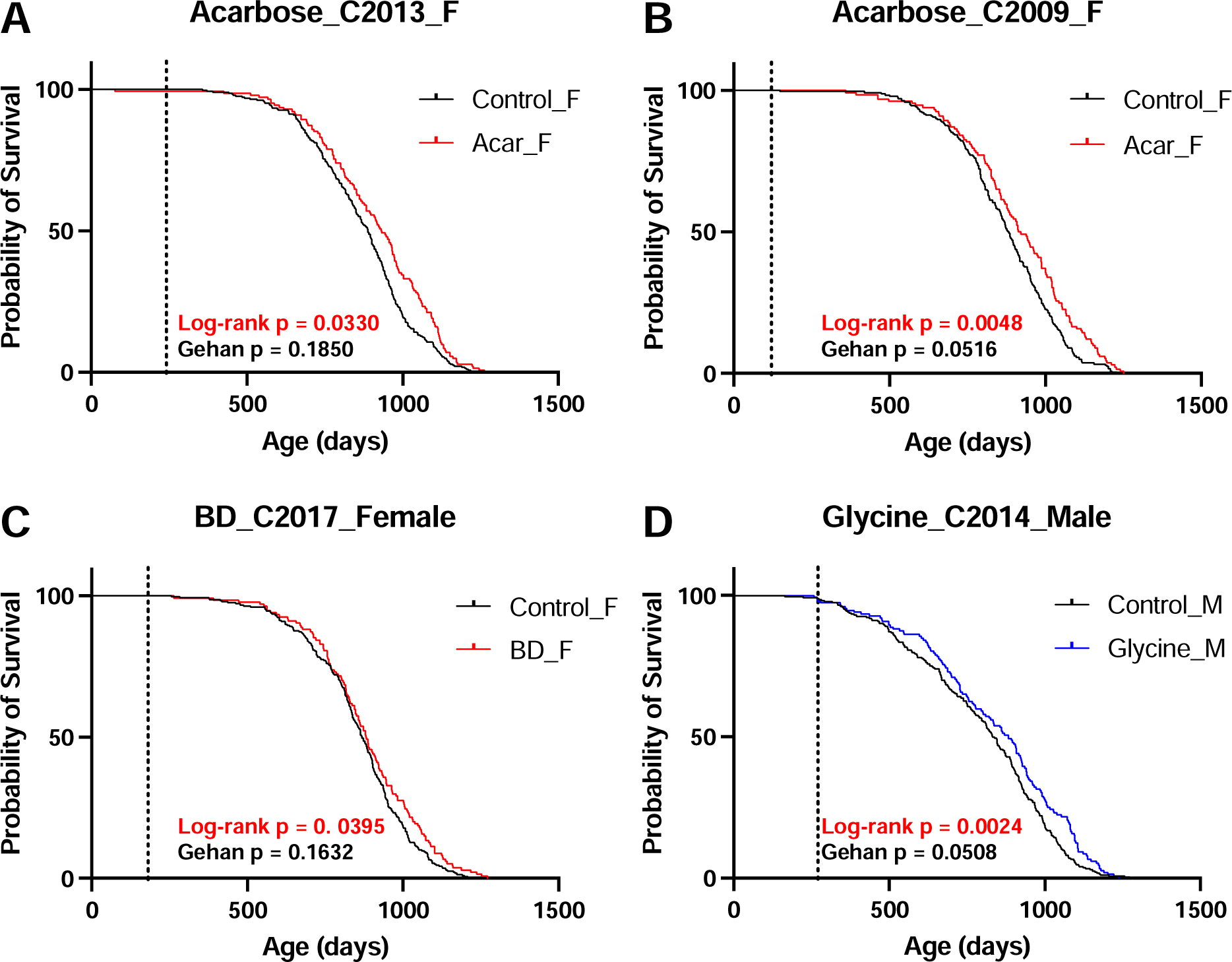
Survival curves for interventions that significantly increased survival according to the log-rank test but lost significance in the Gehan test. **(A)** Acarbose treated (400 ppm, from 8 months, n = 139) and control (n = 287) female mice. **(B)** Acarbose treated (1000 ppm, from 4 months, n = 132) and control (n = 242) females. **(C)** (R/S)-1,3-butanediol treated (100000 ppm, from 6 months, n = 134) and control (n = 276) females. **(D)** Glycine treated (80000 ppm, from 9 months, n = 153) and control (n = 273) males.

**Fig. S3.**
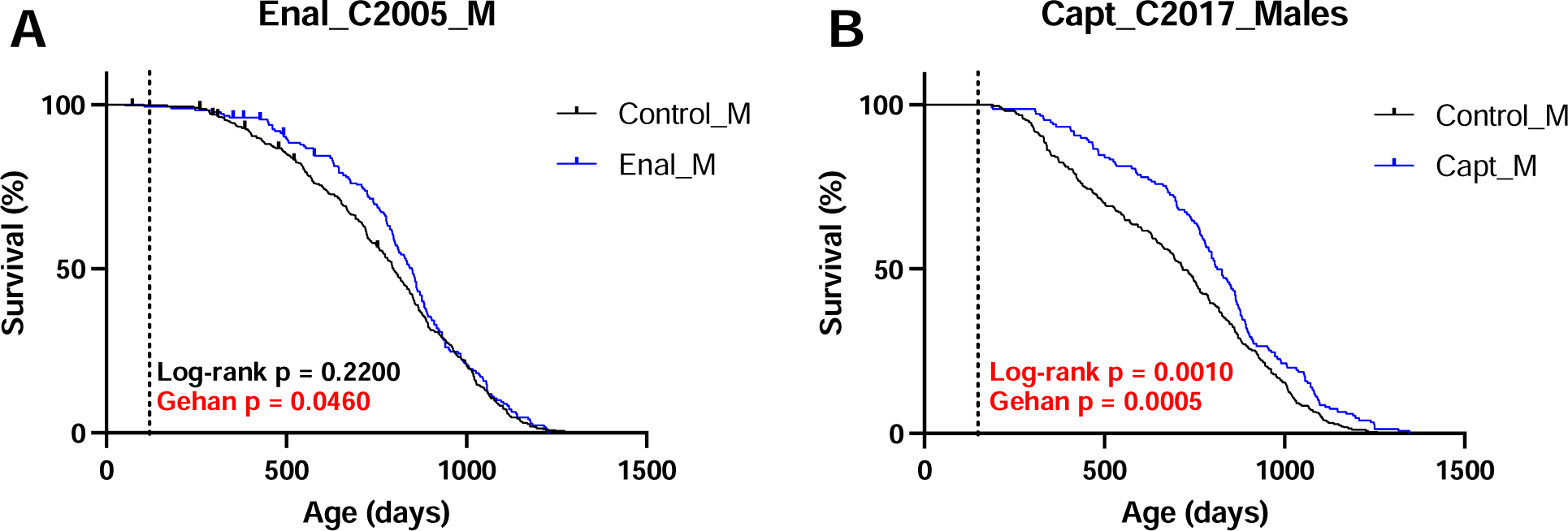
**Survival curves for (A)** Enalapril treated (120 ppm, from 4-month-old, n = 170) and control (n = 357) male mice, and **(B)** Captopril treated (180 ppm, from 5-month-old, n = 150) and control (n = 285) male mice.

